# Exploring the Conformational Transition Between the Fully Folded and Locally Unfolded Substates of the *Escherichia coli* thiol peroxidase

**DOI:** 10.1101/2019.12.26.888669

**Authors:** Diego S. Vazquez, Ari Zeida, William A. Agudelo, Mónica Montes, Gerardo Ferrer-Sueta, Javier Santos

**Affiliations:** Laboratorio de Expresión y Plegado de Proteínas, Departamento de Ciencia y Tecnología, Universidad Nacional de Quilmes, Roque Sáenz Peña 352, Bernal, Buenos Aires, Argentina; Instituto Multidisciplinario de Biología Celular (IMBICE), Calle 526 y Camino General Belgrano, La Plata, Buenos Aires, Argentina; Departamento de Bioquímica y Centro de Investigaciones Biomédicas (CEINBIO), Facultad de Medicina, Universidad de la República, Avda. General Flores 2125, Montevideo, Uruguay; Fundación Instituto de Inmunología de Colombia (FIDIC), Avda. 50 N° 26-20, Bogotá D.C., Colombia; Instituto de Química y Fisicoquímica Biológicas (IQUIFIB) “*Prof. Dr. Alejandro C. Paladini*”, Universidad de Buenos Aires and CONICET, Junín 956, Ciudad Autónoma de Buenos Aires, Argentina; Laboratorio de Fisicoquímica Biológica, Instituto de Química Biológica and CEINBIO, Facultad de Ciencias, Universidad de la República, Iguá 4225, Montevideo, Uruguay; Laboratorio de Genómica e Ingeniería de Sistemas Biológicos. Instituto de Biociencias, Biotecnología y Biología Traslacional (iB^3^). Departamento de Fisiología y Biología Molecular y Celular, Facultad de Ciencias Exactas y Naturales, Universidad de Buenos Aires, Intendente Güiraldes 2160, Ciudad Autónoma de Buenos Aires, Argentina; Consejo Nacional de Investigaciones Científicas y Técnicas, Av. Rivadavia 1917 (C1033AAJ), Ciudad Autónoma de Buenos Aires, Argentina

**Keywords:** peroxiredoxin family, conformational landscape, protein dynamics, native ensemble, molecular dynamics simulation, accelerated molecular dynamics

## Abstract

Thiol peroxidase from *Escherichia coli* (*Ec*TPx) is a peroxiredoxin that catalyzes the reduction of different hydroperoxides. During the catalytic cycle of *Ec*TPx, the peroxidatic cysteine (C_P_) is oxidized to a sulfenic acid by peroxide, then the resolving cysteine (C_R_) condenses with the sulfenic acid of C_P_ to form a disulfide bond, which is finally reduced by thioredoxin. Purified *Ec*TPx as dithiol and disulfide behaves as a monomer in close to physiological conditions. Although secondary structure rearrangements are present when comparing different redox states of the enzyme, no significant differences in unfolding free energies are observed under reducing and oxidizing conditions. A conformational change denominated *fully folded (FF) to locally unfolded (LU) transition*, involving a partial unfolding of αH2 and αH3 helices, must occur to enable the formation of the disulfide bond since the catalytic cysteines are 12 Å apart in the FF conformation of *Ec*TPx. To explore this crucial process, the mechanism of the FF→LU and the LU→FF transitions were studied using long time scale conventional molecular dynamic simulations and an enhanced conformational sampling technique for different oxidation and protonation states of C_P_ and/or C_R_. Our results suggest that the FF→LU transition has a higher associated energy barrier than the refolding LU→FF process in agreement with the relatively slow experimental turnover number of *Ec*TPx. Furthermore, *in silico* designed single-point mutants of the αH3 enhanced locally unfolding events, suggesting that the native FF interactions in the active site are not evolutionary optimized to fully speed-up the conformational transition of wild-type *Ec*TPx.

## Introduction

*Escherichia coli* thiol peroxidase (*Ec*TPx, UniProt entry P0A862) is a small and globular protein of 167 residues belonging to the peroxiredoxin (Prx) family [1–4]. *Ec*TPx shares the prototypical thioredoxin fold, characterized by a seven strand β sheet core surrounded by five helices with an N-terminal β hairpin extension (Figure 1A). Members of the TPx subfamily, originally named p20 or scavengases [5,6], were identified as bacterial atypical 2-Cys Prxs, as both catalytic cysteines (Cys) are located in the same polypeptide molecule, and conform one of the less phylogenetically diverse subfamilies known [7].

**Figure 1:**
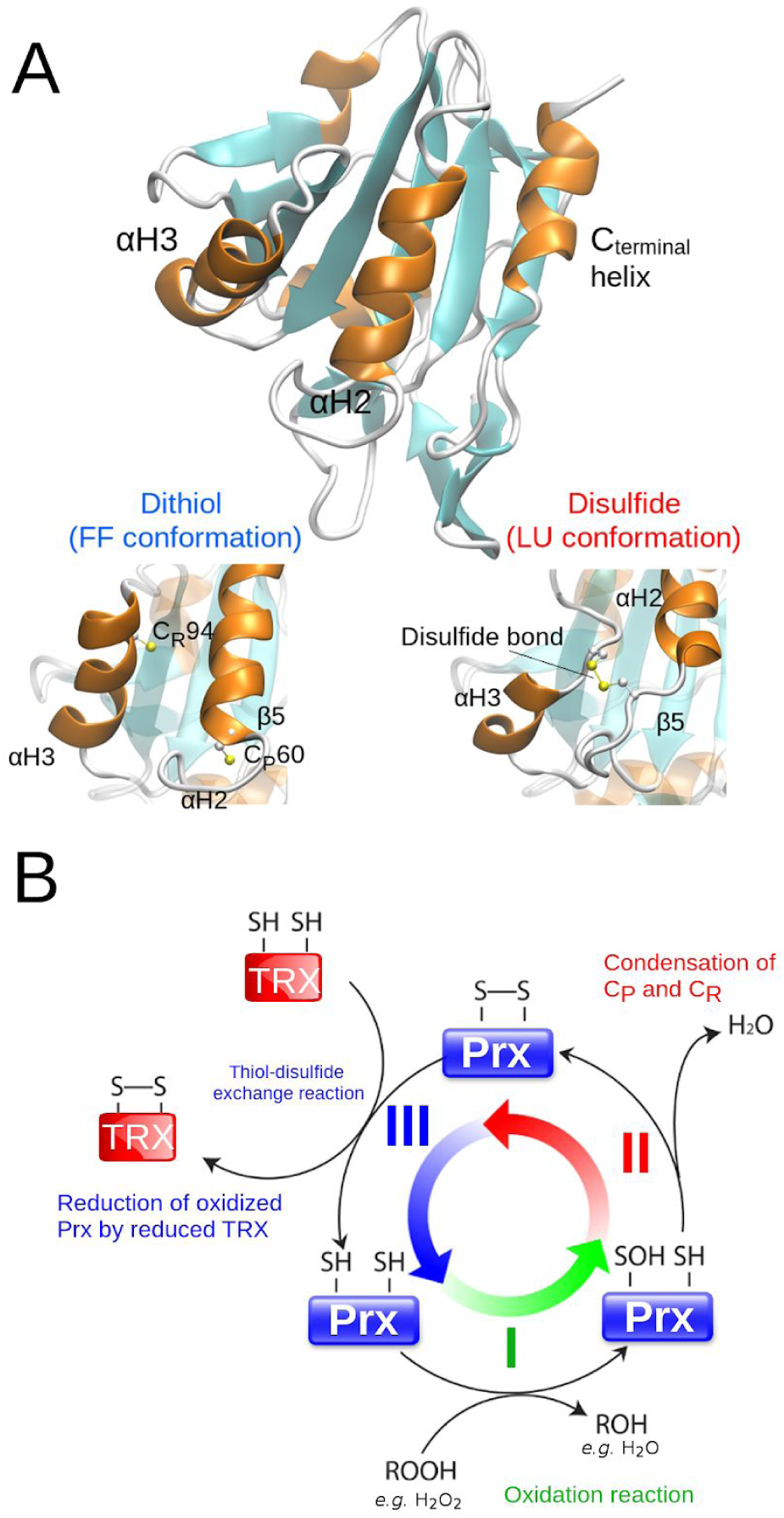
EcTPx X-ray Structure and Catalytic Mechanism. Cartoon representation of the *Ec*TPx structure constituted by a laminar seven strand β sheet core surrounded by five helices (Panel A). The FF (reduced state, PDB ID 3HVV) and LU conformations (oxidized state, PDB ID 3HVS chain A) are shown below. The C_P_ and C_R_ are highlighted. The general three-step catalytic cycle for atypical 2-Cys Prxs is shown in panel B and involves: I) the peroxidation of C_P_ to form the sulfenic acid intermediate (C_P_-SOH); II) the formation of the intramolecular disulfide bond between C_P_ and C_R_, and; III) the reduction of the disulfide by a thiol oxidoreductase (TRX).

*Ec*TPx reduces H_2_O_2_ [8] and alkyl hydroperoxides [9] via a reactive and absolutely conserved peroxidatic Cys (C_P_) surrounded by the highly conserved PxxxT/SxxC_P_ motif [10] and an Arg residue that constitute the active site [11]. The first step of the catalytic mechanism involves the nucleophilic attack of the C_P_ sulfur atom on the peroxide yielding the sulfenic acid intermediate (Cys-SOH) and a water molecule if the substrate is H_2_O_2_. The progress in the catalytic cycle depends on the presence and location of a second Cys residue (resolving Cys or C_R_), that react with the sulfenic acid yielding a disulfide bond (R-C_P_-S-S-C_R_-R), releasing a water molecule as a leaving group. Finally, the disulfide bond is reduced by thioredoxin (TRX), through a typical thiol-disulfide exchange reaction (Figure 1B, [12]).

The catalytic cycle goes through at least one major conformational transition, the so-called *fully folded* (FF) to *locally unfolded* (LU) transition [2,13–15], as C_P_ (Cys60) and C_R_ (Cys94) are more than 12 Å apart in the crystallographic structure of the reduced (dithiol) state (Figure 1A, [13]). Whether the conformational equilibrium is modulated by the redox and protonation state of C_P_ and/or C_R_ is still an open question. The transition involves a partial unfolding of the C-terminal region of αH3 (residues 86 to 97), together with a smaller local unfolding of the N-terminal region of αH2 and the preceding loop connected to the β5 strand and changes in the solvent accessibility of C_P_ (Figure 1A).

The specificity constant of *Ec*TPx for H_2_O_2_ (*k*_cat_/K_M_ = 8.9 × 10^4^ M^−1^ s^−1^, [16]) is in the range observed for other TPx-Prx subfamily members [17]. The kinetic parameters for the individual steps in the reduction of H_2_O_2_ by TPx (catalyzed by TRX) have been reported [8], *k*_cat_= 76 s^−1^ (H_2_O_2_), K_M_ = 1.7 mM (H_2_O_2_) and K_M_ = 25.5 µM (TRX), and indicate that the most likely rate-limiting step of the cycle is the peroxidation (Reaction I, Figure 1) since large concentrations of H_2_O_2_ would be needed to saturate the enzyme. The only other hydroperoxide tested, cumene hydroperoxide, has a much smaller K_M_ (9.1 µM) pointing to a specialization of the enzyme for larger hydrophobic peroxides. In those cases, reduction and/or resolution reactions may become rate-limiting and both reactions involve the conformational transitions under study, FF→LU and LU→FF, respectively. Crystallographic and kinetic approaches have provided evidence describing the FF and LU conformations in different Prx subfamilies [8,13,18,19]. However, a thorough dynamic characterization of the FF→LU and LU→FF transitions is not available yet. To shed light on this process, the mechanism of *Ec*TPx conformational transitions with C_P_ modeled in the thiol, thiolate, sulfenic acid and sulfenate states and C_R_ modeled in the thiol and thiolate states were studied in the present work, using long time scale conventional molecular dynamic (cMD) simulations to study the intrinsic dynamics of the FF and LU conformational substates together with the enhanced sampling technique accelerated molecular dynamics (aMD) to describe conformational changes. Additionally, a set of biophysical experiments were carried out in order to characterize the oligomeric state, the secondary/tertiary structure content and the thermodynamic properties of the dithiol and disulfide states of the enzyme.

## Results

### Expression, Purification and Experimental Characterization of Wild-type *Ec*TPx

*Escherichia coli* TPx was overexpressed as a soluble protein and purified in the oxidized form (disulfide bond between C_P_60 and C_R_94), as judged by the absence of free thiols in the native state (data not shown). The molecular weight (MW) was evaluated by ES-MS (17701 Da) and coincides within 1 Da with the expected mass deduced from protein sequence, considering the loss of two protons in the oxidized state and the lack of the initial Met residue. The secondary and tertiary structure content of *Ec*TPx in the dithiol (preincubated with 1 mM DTT) and disulfide states were evaluated by circular dichroism (CD) spectroscopy. The spectrum in the far-UV region show native-like signatures compatible with a folded protein (Figure 2A). The lower negative signal at 222 nm and lower positive signal at 195 nm of the far-UV CD spectrum in the oxidized form account for the loss of periodic helical structure compatible with the locally unfolded (LU) conformation. Furthermore, the near-UV CD spectrum shows a significant gain of negative band in the 270-285 nm region, in oxidizing conditions, which is also compatible with the formation of a disulfide bond (Figure 2B), reflecting a change in the asymmetric environment of aromatic residues near the active site (Figure 1B).

**Figure 2:**
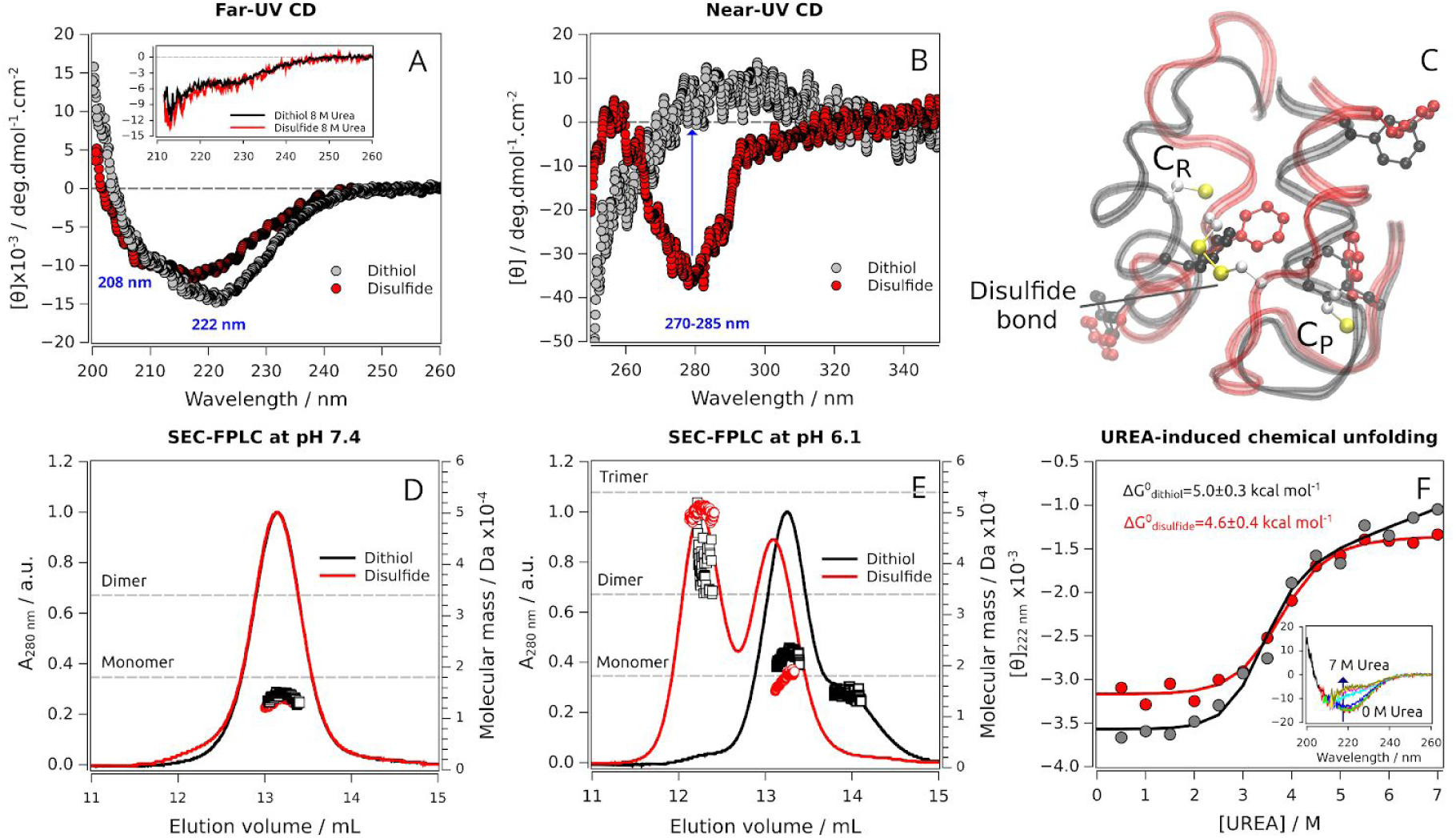
Biophysical Characterization of Wild-type EcTPx. The far-UV (A) and near-UV CD spectra (B) are shown for the reduced (black) and oxidized (red) states. Additionally, the spectra for each oxidation state in the presence of 8.0 M GdmCl are shown in the inset. Aromatic residues near the active site in the dithiol (black) and disulfide (red) states area highlighted in panel C. The SEC-FPLC elution profile and the oligomeric state was studied by static light scattering at pH 7.4 (D) and pH 6.1 (E) in reduced (black line) and oxidized (red line) conditions, respectively. The theoretical mass of the monomer (17.7 kDa), homodimer (35.4 kDa) and homotrimer (53.1 kDa) are represented by dashed grey lines. The urea-induced chemical unfolding of the dithiol and disulfide states of wild-type *Ec*TPx is shown in panel F. Far-UV CD spectra of reduced *Ec*TPX at different urea concentrations are shown in the inset.

A previous bioinformatic and structural analysis reported that the TPx subfamily shows a region of highly conserved hydrophobic residues that in the oxidized state forms an intermolecular interface involving a surface area of ∼1400 Å^2^ [8], suggesting a role for the stabilization of the dimeric form of *Ec*TPx in solution. In the same fashion, experimental measurements carried out by analytical ultracentrifugation showed that the dimer is the predominant species in solution above 40 *μ*M. Furthermore, the authors concluded that the dimer interface is involved in the stabilization of the active site [8]. To evaluate the oligomeric state of *Ec*TPx at low micromolar concentrations (10-20 *μ*M), we performed SEC-FPLC experiments monitored by means of a multiangle light scattering detector at neutral pH and moderate ionic strength (20 mM Tris-HCl, 100 mM NaCl, pH 7.4). The predominant species in both reduced and oxidized conditions exhibited molecular masses compatible with a monomer species (Figure 2D). These observations are also in agreement with SEC experiments carried out with a TPx from *Helicobacter pylori* [20] and the PrxQ from *Xanthomonas campestris* [21] that shows a predominant monomeric species in both reduced and oxidized conditions. To test the effect of ionic strength in the monomer-dimer equilibrium, SEC-FPLC experiments were carried out increasing the NaCl concentration up to 400 mM. No dimeric species were observed, neither in reduced nor oxidized conditions (data not shown).

Curiously, the oxidized enzyme at pH 6.1 forms a homotrimer with an average molecular mass of 49.9 ± 1.1 kDa (Figure 2E). On the other hand, only a much less abundant and highly-dispersed population, compatible with dimeric or trimeric forms, was detected for the reduced state. Even more, a species with a molecular mass compatible with the monomer was observed with a slightly higher elution volume, compared to that observed at pH 7.4 (Figure 2D and E), possibly due to a particular interaction of the protein with the Superose-12 matrix at this pH.

The urea-induced chemical unfolding of the reduced and oxidized states of *Ec*TPx are cooperative, indicating that both states are organized and contain a native protein core. The difference in free energy between the unfolded and folded state obtained by equilibrium unfolding experiments are nearly unchanged in the oxidized or reduced states of *Ec*TPx (Figure 2F, ΔG°_NU_< 0.5 kcal mol^−1^) which is in agreement with the low difference in the melting temperature (*T*_m_) observed for the TPx from *H. pylori* [20] in the reduced (*T*_m_ = 66.6 °C) and oxidized conditions (*T*_m_ = 65.0 °C). However, it is worthy to note that the unfolded state of the dithiol and disulfide states of *Ec*TPx is structurally different because of the presence of the disulfide bond, making difficult the comparison between them in energetics terms.

### Structural Characterization of the Native FF and LU Conformational Substates by MD Simulations

To get insights about the conformational transitions and internal motions that take place during the catalytic cycle of *Ec*TPx, we performed molecular dynamics (MD) simulations. Several systems were evaluated: (I) the wild-type *Ec*TPx substituting Ser60 residue for Cys (C_P_) as thiol, thiolate, sulfenic acid or sulfenate (starting from FF conformation, PDB ID 3HVV); (II) the wild-type *Ec*TPx with C_P_ as thiol, thiolate, sulfenic acid or sulfenate starting from a locally-unfolded-like conformation (LU*, PDB ID 3HVS chain A); (III) the wild-type *Ec*TPx in the oxidized state (disulfide bond, PDB ID 3HVS chain A); (IV) as system II but evaluating the four possible protonation combinations for both Cys (C_P_ and C_R_); (V) three single-point mutants of the aH3 helix, starting from FF conformation, with C_P_ and C_R_ modeled in the sulfenic acid and thiol form, respectively, substituting the residues Arg92, Phe93 or Glu97 of αH3 helix by Gly (see text below).

In order to validate our computational simulation setup, a comparative analysis was performed between the root-mean-square deviation (RMSD) of the experimental structures of *Bacillus subtilis* TPx (*Bs*TPx) determined by NMR [19] in the truly reduced state (dithiol) with the B factor of the crystallographic structures of *Ec*TPx [8] and the root-mean-square fluctuation (RMSF) of our conventional MD (cMD) simulations in the dithiol and disulfide states (Figure 3). Remarkably, our cMD simulations of *Ec*TPx captures the fundamental conformational features of the reduced and oxidized ensembles. The RMSF profiles found in our simulations reproduce very well the ensemble heterogeneities inferred from B factors and RMSD profiles corresponding to *Ec*Tpx and *Bs*TPx, respectively.

**Figure 3:**
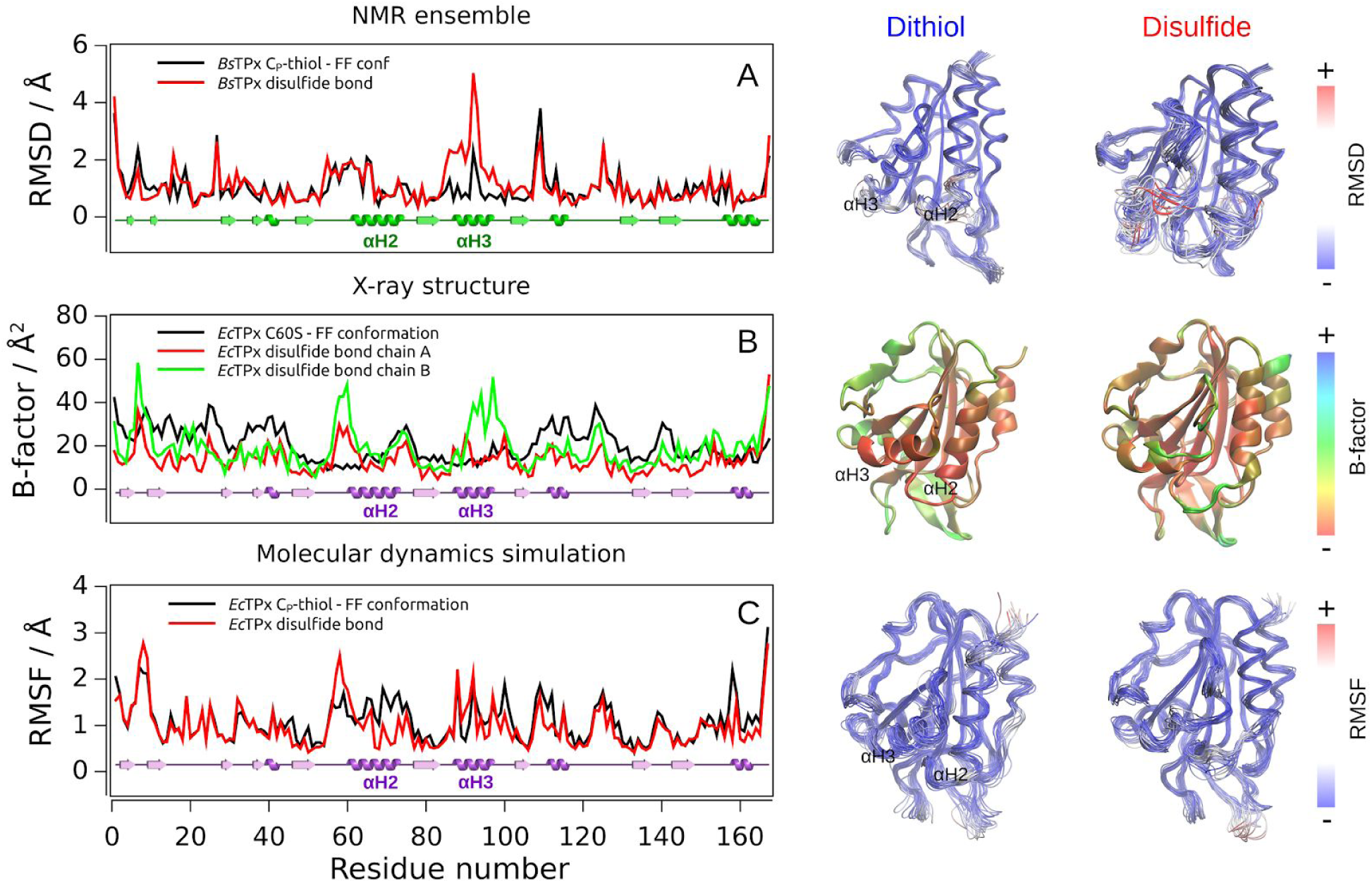
Conformational Heterogeneity of E. coli and B. subtilis TPx. The root-mean-square deviation *per* residue (RMSD) of the 20 lowest-energy structures of *Bs*TPx determined by NMR in the reduced state (PDB ID 2JSZ, black line) and the disulfide bond state (PDB ID 2JSY, red line) are shown in panel A. The B-factor of *Ec*TPx in the reduced state (PDB ID 3HVV, black line) and disulfide bond state (PDB ID 3HVS chains A and B, red and green lines, respectively) are shown in panel B. The root-mean-square fluctuations (RMSF) of the cMD simulations of both redox states of *Ec*TPx are shown in panel C with C_P_ and C_R_ modeled as thiols. The values are represented in the corresponding 3D structure in the right panel. The linear secondary structure and the αH2 and αH3 regions are indicated for clarity.

These analyses pointed out the possibility to explore the dynamics of the FF and LU ensembles for the physiological and relevant chemical states of C_P_: thiol, thiolate, sulfenic acid and sulfenate species. In this regard, there is a clear difference in the interactions that each species can form and how they may influence in the FF ⇆ LU transition. The thiolate species is a hydrogen bond (HB) acceptor and can establish salt bridge interactions due to its negative net charge, so it is expected to stabilize the FF conformation through interactions with polar and charged residues in the vicinity, particularly the conserved Thr57 and Arg132 of the active site. On the other hand, the thiol group is a poor HB donor due to the low electronegativity and large size of the sulfur atom, moreover, it cannot establish ionic interactions. Once the C_P_ is oxidized, the presence of the C_P_-SOH group, that can form stronger HB due to the extra oxygen atom, could affect the HB network of the active site. Additionally, being more voluminous than thiol or thiolate, the sulfenic acid species may affect the conformational architecture of the active site [22]. Furthermore, if the sulfenate is the predominant species at neutral pH [23], the oxygen can act also as an HB acceptor.

We clusterized different structures obtained in the cMDs using principal component analysis (PCA) as clustering criteria and identify key residues involved in large conformational movements. For reduced and oxidized ensembles, the first two eigenvectors fall near the origin of the Cartesian axis, suggesting that the systems are sampling conformations close to the ones observed in the crystallographic structures. Moreover, the overlap of the root-mean-square inner product (RMSIP) value of 0.4, denotes a low similarity of the overall dynamics [24]. It is worthy to note that the essential subspace in each conformational substate is expected to change due to the conformational restriction imparted by the disulfide bond between C_P_ and C_R_.

The deprotonated C_P_-thiolate and C_P_-sulfenate species keep nearly unchanged with respect to the starting structure, establishing ionic interactions with Thr57 and Arg132. Although the protonated forms transiently got closer between the catalytic Cys residues, in any case, local unfolding transitions were observed, as judged by the secondary structure analysis (SSC, data not shown). This observation suggests that the thiolate species, that is the most abundant form under physiological conditions and reactive one, contributes to the stability of FF conformations.

For the disulfide bond formation, two things must occur: the approximation of catalytic Cys residues and also the loss of secondary structure, with a concomitant increment in local fluctuation. When cMD simulations started from the LU conformation, the deprotonated species, particularly the C_P_-thiolate system, present a maximum peak with a C_P_-C_R_ distance of 5.6 ± 0.3 Å, close to the starting distance at the beginning of the simulation (Figure 4, right panel). Nevertheless, fully refolding events compatible with the FF conformation was not observed in any case. These results are interesting because, although the form that reacts with C_R_ is the sulfenic acid form of C_P_, the sulfenate species would remain closer to the C_R_, transiently stabilizing the LU conformation, necessary for the reaction. Additionally, it has been found that although the preferent reaction is between C_P_ sulfenic and C_R_ thiolate, the reaction with the C_P_ sulfenate also occurs [23].

**Figure 4:**
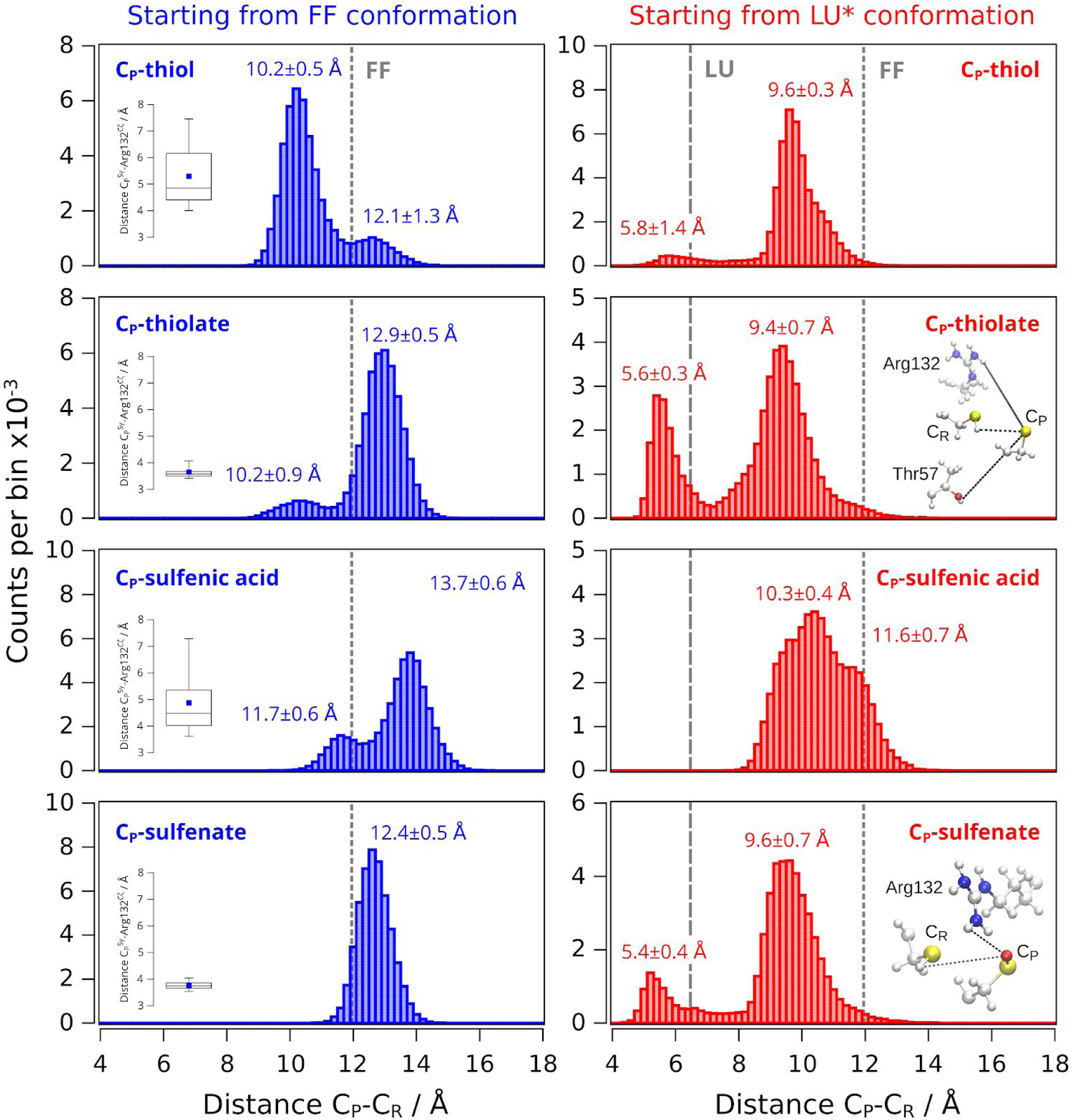
Analysis of C_P_-C_R_ Distance Distributions. Histogram The C_P_-C_R_ distance of the Cα atoms through the cMD simulations starting from the FF (left panel) or LU* (right panel) conformations for different oxidation and protonation states of C_P_. The distributions of the distance between the sulfur atom of C_P_ and the C**ζ** atom of Arg132 are shown as boxplot insets. The C_R_ was modeled as a thiol in all systems. Vertical grey lines indicate the distance values between Cα atom in the corresponding crystallographic conformation. The values reported corresponds to the mean ± 1σ of the multimodal Gaussian fit.

### Exploring the FF→LU Conformational Transition by Accelerated MD Simulations

Helix-coil transitions usually involve surpassing high energy barriers, thus cMD simulations become insufficient to study this kind of conformational rearrangements. To overcome this obstacle, we employed accelerated molecular dynamics (aMD) simulations, an enhanced sampling technique that extends the effective simulation time scale to the order of microseconds [25]. Due to the unknown value of the energy barrier involved in the FF→LU process, four multiples λ values of the acceleration factor α were applied to the *Ec*TPx C_P_-sulfenic acid in the FF conformation system ranging from 0.3 to 4.0 collecting a total 1.75×10^9^ simulation steps, equivalent to a computational cost of 3.5 μs. Although at λ = 0.3 the system shows a more spread out map along the first two eigenvectors compared with the cMD simulation (Figure 5A), the conformations sampled in the cluster of maximal frequency are still similar to those sampled by the cMD simulation, as pointed out by the low value of the average ΔRMSD of 0.5 Å, a RMSIP value of 0.7 and a C_P_-C_R_ distance of 12.2 ± 0.8 Å between Cα atoms. Interestingly, multiple solvent exposure events of the C_P_ residue were observed at λ = 1.0 characterized by a ΔSASA of 88.2 ± 20.2 Å^2^. Nevertheless, this was not accompanied by a significant local unfolding of the *α*H2 neither an approach of the catalytic Cys which keep at an average distance of 12.1 ± 0.7 Å (Figure 5B). This fact could indicate that the Cys approximation and the concomitant condensation of the disulfide bond, involves two independent stages where the unfolding of one of the helices does not induce the unfolding of the other. Similar results were observed at λ = 2.0 where the C_P_-C_R_ distance remained stable at 11.9 ± 0.6 Å without evidencing significant changes in the SSC of αH2 or αH3 (Figure 5C).

**Figure 5:**
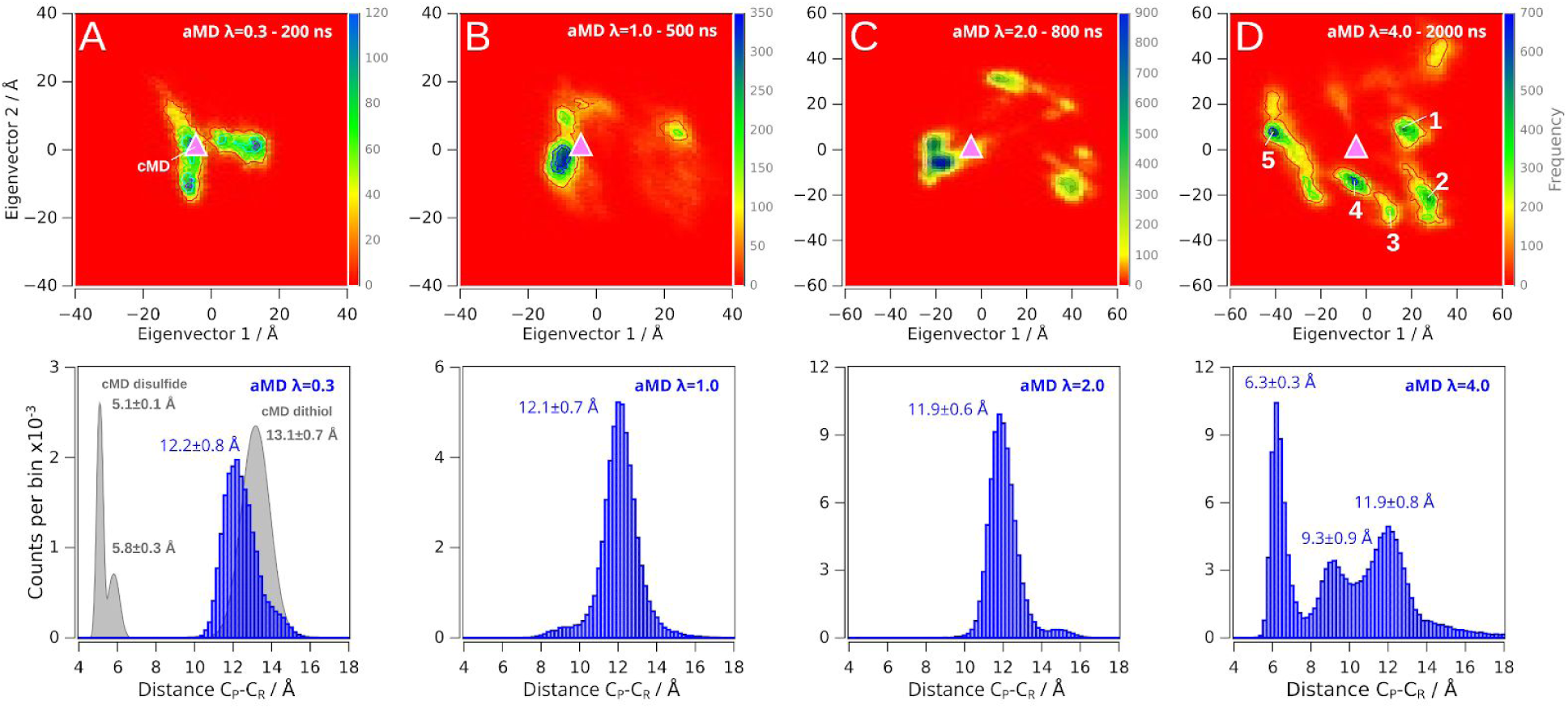
Effect of λ Factor on the Conformational Sampling of Wild-type EcTPx. Bidimensional heatmaps of the eigenvectors 1 and 2 projected onto the aMD trajectory for *λ* factor values of 0.3 (A), 1.0 (B), 2.0 (C), and 4.0 (D) are shown in the top. The pink triangle denotes the position of the maximum frequency in the cMD simulation starting from the FF conformational substate with C_P_ and C_R_ modeled as the sulfenic acid and thiol, respectively. The numbers in panel D denotes the wells analyzed in the main text. The corresponding histogram of the C_P_-C_R_ distance for Cα atoms through all the trajectory is shown below each panel. The distributions for the dithiol and disulfide states of the cMD simulations are shown as a reference (grey). The values reported corresponds to the mean ± 1σ of the multimodal Gaussian fit.

It is worth to note that an increase in λ parameter (*i.e.* a decrease in the acceleration factor α) will result in a decrease in ΔV(*r*) (the boost potential) and then, a longer simulation time will be needed to properly sample the protein conformational space [26]. For this reason, a microsecond time scale simulation was performed at λ = 4.0. The bidimensional heatmap shows more populated minima besides the crystallographic basin (Figure 5D). To characterize the conformations accumulated in each energy minimum, a clustering analysis over the projection of the covariance matrix in the accelerated simulation was conducted. Five minima clusters preserving native and non-native signatures were found at λ = 4.0 and four of them are characterized to hold conformations which are far away from the crystallographic FF conformational substate. The conformations clustered in the well 1 have a slight shift in the position of αH2 and a twist of 40° in the αH3 refereed to the cMD simulation with no significant loss of helicity. For wells 2 to 5, a drastic sheet-to-helix transition involved the antiparallel β3-β4 (residues 22 to 44) to an α-helix was formed. The loss of the β3-β4 interactions affects the local dynamics of β7 which is in close proximity with the αH3 helix. Nevertheless, a similar behaviour was observed for αH2 and αH3 than in the well 1. Well 3 holds structures in which the helix observed in the wells 2-5 breaks into 2 parallel α-helix with a partial unfolding of αH3 and interestingly near 70 % of the conformations have the catalytic Cys at less than 4 Å. Surprisingly, a complete loss of secondary structure of the αH3 helix was found in well 4 but the average C_P_-C_R_ distance remained at 12.1 ± 0.8 Å and only 5 % of the conformers the catalytic Cys approach to less than 4 Å, reinforcing the idea about the independence between C_P_-C_R_ approximation and local unfolding phenomena.

### Exploring the LU→FF Conformational Transition by Accelerated MD Simulations

To explore the conformational transition from the LU to the FF conformation, we prepared four additional systems using a LU-like (LU*) conformation as starting structures where the C_P_ and C_R_ (in the reduced state) were combined in the protonated (P) or deprotonated (D) states (I: C_P_^P^ and C_R_^P^; II: C_P_^P^ and C_R_^D^; III: C_P_^D^ and C_R_^P^; and IV: C_P_^D^ and C_R_^D^). The systems were subjected to 100 ns of cMD simulation using a distance restraint of 3.5 Å between the Cα atoms of C_P_ and C_R_. Then four aMD simulation replicas (7.5×10^7^ simulation steps *per* replica, equivalent to 150 ns) were performed for each system using the last structure as initial coordinates for the unrestricted simulations assigning new random velocities.

The analysis of the simulation of the C_P_^P^-C_R_^P^ system showed an ensemble population with a local RMSD of 2.2 Å and a C_P_-C_R_ distance distribution of 12.4 ± 0.8 Å between Cα atoms (Figure 6). The analysis of the helical content *per* residue (Figure 7A) through the simulations, revealed that the protonation of C_R_ rises the helical propensity of the first segment of αH3 comprising the residues 87 to 90, to more than 90 % on the whole trajectory. Furthermore, when the simulations were analyzed in three sections: the first part (0-50 ns), seconds part (50-100 ns), and the third part (100-150 ns) we observed that during the last part of the simulations (100-150 ns) the helical structure of the complete αH3 (residues 87-96) was acquired (Figure 7A). On the other hand, when both Cys residues were modeled as thiolate (system IV, C_P_^D^ and C_R_^D^), the stabilization of helical structure for the first seven residues of αH3 (residues 87-93) was significantly reduced. Intermediate behaviors were observed for systems II and III which have one Cys as thiolate and the other as thiol (C_P_^P^-C_R_^D^ and C_P_^D^-C_R_^P^, Figures 6 and 7).

**Figure 6:**
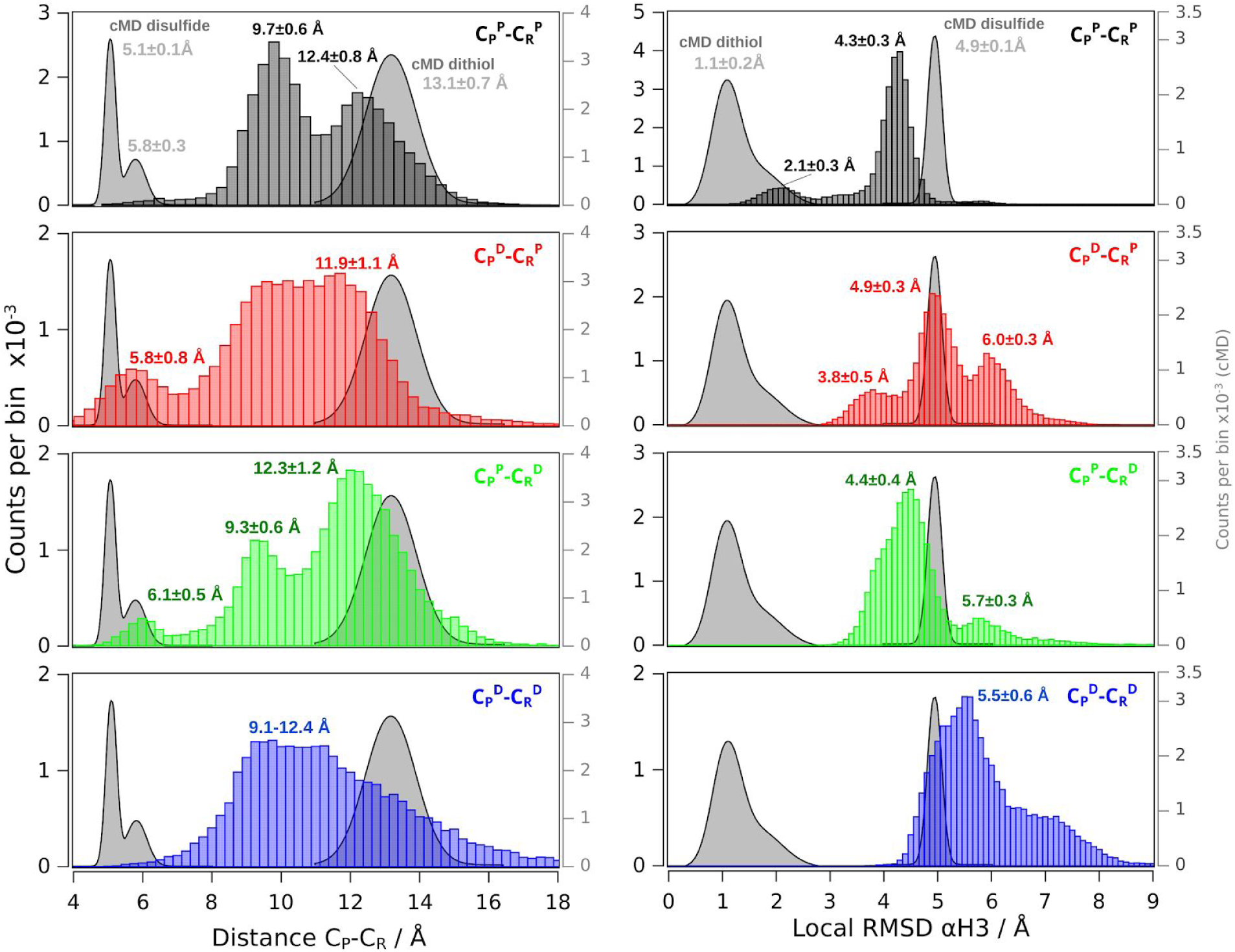
Effect of Protonation State of C_P_ and C_R_ on the Acquisition of the LU→FF Conformational Transition. Distribution of local RMSD of the αH3 was performed over the 150 ns aMD simulation for the C_P_^P^-C_R_^P^ (black), C_P_^P^-C_R_^D^ (red), C_P_^D^-C_R_^P^ (green) and C_P_^D^-C_R_^D^ (blue) systems are shown in the right panel. The crystallographic FF conformation (PDB ID 3HVV) was used for all systems as reference for global fitting and subsequent local RMSD calculation of the αH3. The histograms of the C_P_ and C_R_ distance between the Cα atoms are shown in the left panel. The distance distribution for the FF and LU systems in the cMD simulations is shown as a reference (grey). The values reported corresponds to the mean ± 1σ of the multimodal Gaussian fit. The superscript P and D labels denote the protonated or deprotonated state of the catalytic Cys, respectively.

**Figure 7:**
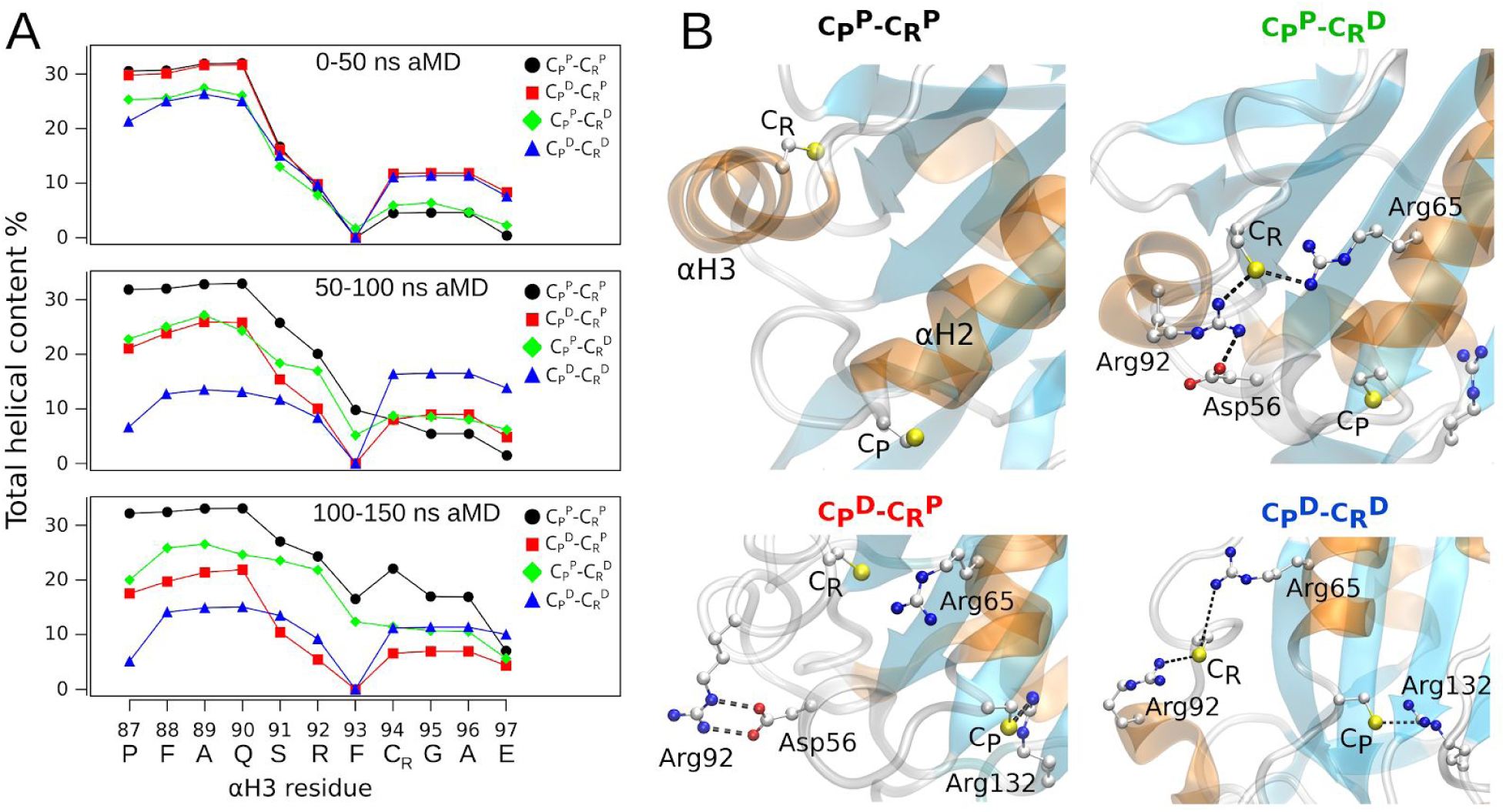
Total Helical Content of the αH3. The percentage helical content of αH3 for the C_P_^P^-C_R_^P^ (black circles), C_P_^P^-C_R_^D^ (red squares), C_P_^D^-C_R_^P^ (green diamonds) and C_P_^D^-C_R_^D^ (blue up triangles) systems analyzed between 0-50, 50-100 and 100-150 ns of each aMD simulation are shown in panel A. The primary sequence of αH3 is shown in the *x*-axis. Representative snapshots of each system are shown in panel B. The superscript labels P and D correspond to the protonated or deprotonated state of the catalytic Cys, respectively.

Deprotonated Cys residues established ionic pair interactions with Arg92, Arg65 (in the case of C_P_) and Arg132 (in the case of C_R_, Figure 7B). These interactions presumably impede the helical structure acquisition, or at least delay its induction, stabilizing the LU conformations and keeping the system trapped, suggesting that the protonation state of the C_P_ residue is a key feature for the modulation of the equilibrium between the LU and FF conformations. Noticeably, Arg65, Arg92 and Arg132 residues are conserved in the TPx subfamilies [1]. This behavior is also in agreement with a high local RMSD of the αH3 helix and broad distribution of the C_P_-C_R_ distance (Figure 6).

### Effects of αH3 Helix Single-point Mutations on the FF Conformational Ensemble

The analysis of the interactions in FF or LU ensembles showed that the FF and LU conformations are stabilized by different hydrogen bond (HB) networks. The segment comprised between residues 86 to 92, involving the loop connecting the β6 and the first half of αH3, is strongly stabilized by HBs in the FF ensemble and hydrophobic interactions of Phe93 side chain with the protein core (Figure 8, left panel). Noticeably, the second half of αH3 helix (residues 93 to 97), containing the C_R_, is stabilized by only one interaction between Glu97 and Arg65. On the other hand, in the LU ensemble, three HB are formed in the same region: two HB between the side chain N^ζ^ of Arg65 with the backbone oxygen of Glu97 and C_P_, and one between the backbone oxygen of Phe93 with the backbone N-amide of Glu97; plus the covalent disulfide bond that globally stabilizes the LU conformations (Figure 8, right panel).

**Figure 8:**
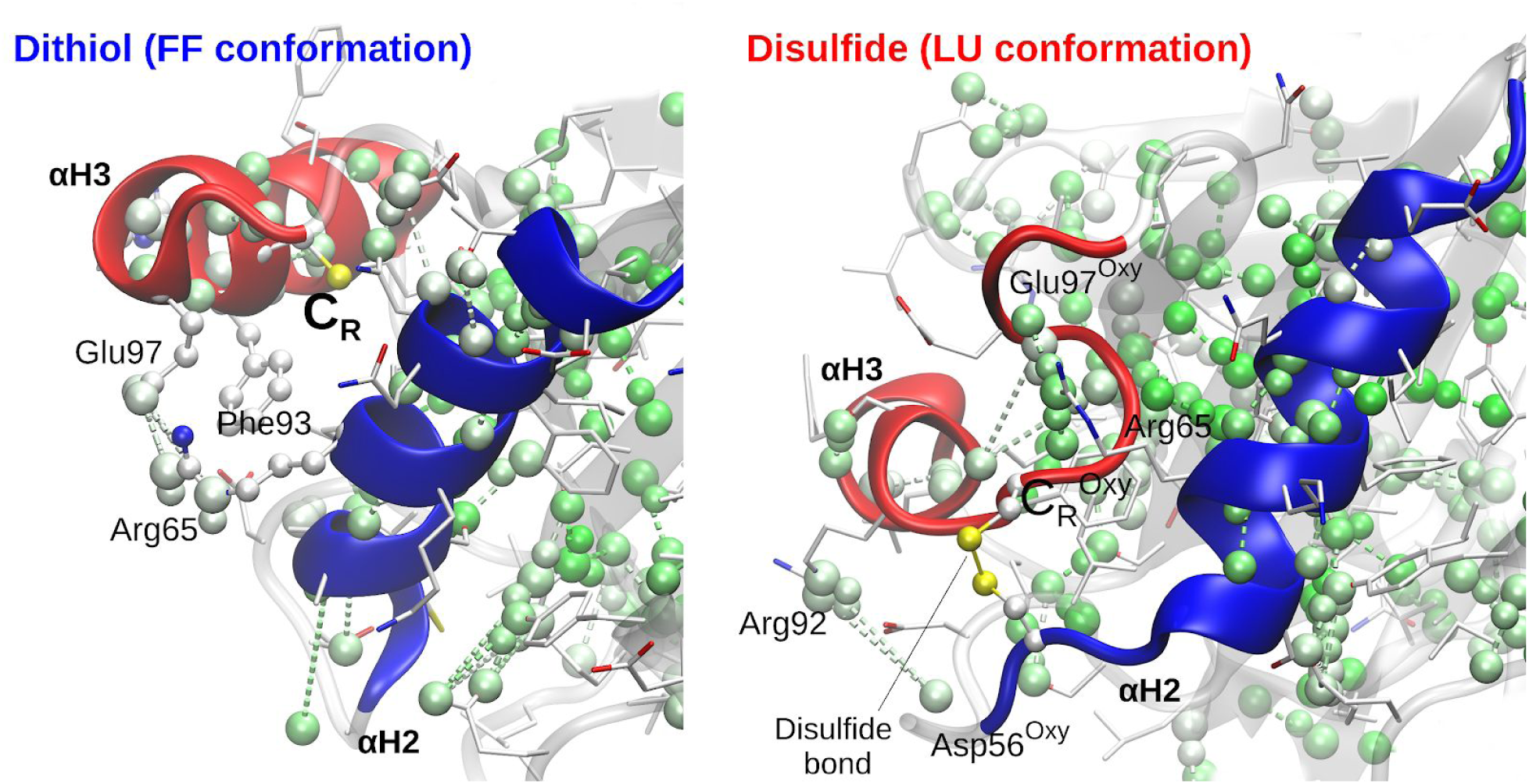
Hydrogen Bond Network Analysis. The HBN was calculated using HBonanza [27] from the cMD trajectories of wild-type *Ec*TPx in the reduced (left panel) and oxidized states (right panel).

To evaluate the effect of altering the HB network and helix propensity, three point mutants were evaluated by MDs, R92G, F93G, and E97G. It is worthy to mention that Gly mutations have two simultaneous effects: first, the truncation of the complete side-chain of the residue and second, the increment of conformational freedom and consequently, the destabilization of helical conformations. The three systems were studied throughout 2 μs cMD simulations. Variants R92G, F93G, and E97G showed RMSD values of 1.3 ± 0.2 Å, 1.6 ± 0.2 Å and 1.9 ± 0.3 Å, respectively. However, significant changes occurred, as inferred by the SSC analysis of αH3 (Figure 9). As expected, wild-type *Ec*TPx and R92G mutation are not perturbed along the simulation either in the SSC of αH3 nor the C_P_-C_R_ distance which kept at 13.9 ± 2.0 Å in concordance to the wild-type variant (average distance C_P_-C_R_ 12.3 ± 2.7 Å, Figure 9A and B). On the contrary, both F93G and E97G mutants show several unfolding transition events of the αH3 helix, concomitantly with the approach of the catalytic Cys to a disulfide bond compatible distance (Figure 9C and D).

**Figure 9:**
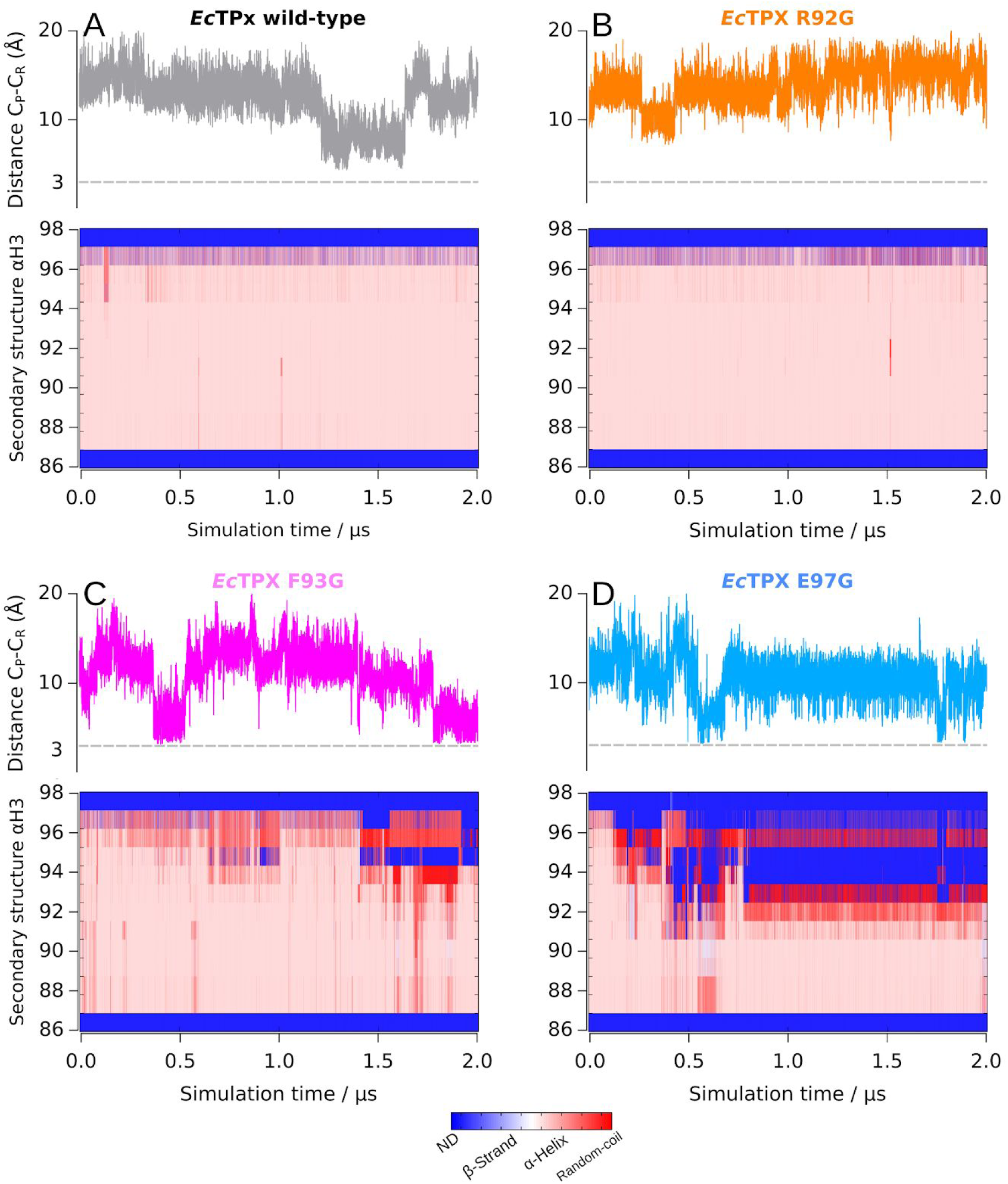
Structural Analysis of EcTPx αH3 Mutants. The C_P_-C_R_ distance together with the SSC calculated by the DSSP algorithm [28] of the αH3 helix are shown for the wild-type variant (A) and the point mutants R92G (B), F93G (C) and E97G (D). The grey dotted line indicates a distance of 3 Å. ND: non-determined.

## Discussion

The study of connections among protein structure, conformational stability, protein dynamics and function is currently an area of intense efforts. The relationships among these key-features define the energy-functional landscape of proteins and comprehend how it occurs is critical to understand enzyme catalysis. Proteins and pathways that include enzymes belonging to thioredoxin (TRX) superfamily have been well established as system models for this kind of studies. Members with very different structural features, mechanisms of action and activities have been studied. In this context, we decided to study experimentally and computationally the enzyme *Ec*TPx, a protein that shares the TRX fold and emerges as an interesting case in the Prx subfamily.

The protein was successfully overexpressed in *E. coli* and purified to homogeneity, and further analyzed under reduced and oxidized conditions in experimental and computational studies. Although the enzyme was previously described as a dimer, as suggested by ultracentrifugation experiments at high concentrations (above 40 µM, [8]), our results demonstrate that, close to physiological conditions and in both redox states, it behaves as a monomer (Figure 2D). The oligomeric features and its influence on the different functions of Prxs is a matter of debate [3,29,30]. Most Prxs behave as homo-oligomers (dimers, decamers, etc.) and changes in the quaternary structure are very dependent on a variety of experimental or cellular conditions [31,32]. Nevertheless, a few reports on members of the TPx and PrxQ subfamilies have already shown by applying different techniques that some Prxs behave as monomers, such as the case of *Bs*TPx [19], PrxQ from *Arabidopsis thaliana* [33] and PrxQ B from *Mycobacterium tuberculosis* [34]. Although key secondary structure rearrangements linked to protein function are present when comparing different redox states of *Ec*TPx, no significant differences in unfolding free energies were observed under reducing and oxidizing conditions. Remarkably, this behavior contrasts with that observed for *Ec*TRX enzyme, the oxidoreductase partner of *Ec*TPx in the catalytic cycle (see Figure 1B). When both redox states of the *Ec*TRX are compared no conformational change are present and only subtle changes in internal motions are observed [16,35,36], whereas the difference in the free energy of unfolding for oxidized state is significantly higher than the corresponding to the reduced state (Δ*G*°_NU, H2O ox_ - Δ*G*°_NU, H2O red_ = 3.5 kcal mol^−1^ at 25 °C, [37,38]). More experiments should be done to understand why such different behaviors were selected through evolution and to elucidate if there is a thermodynamic connection between both proteins.

Structural evidence provided by X-ray crystallography concerning different states of AhpC from *Salmonella typhimurium* initially allowed to infer the existence of complex relationships between conformational exchange and activity, pointing to the existence of at least two different and functionally relevant native substates [20]. In the case of the TPx subfamily, the catalytic reaction comprises a partial local unfolding/refolding event of two non-consecutive alpha-helical elements, bringing the intermediate C_P_-SOH and C_R_ species close enough to react, yielding a disulfide bond and releasing a water molecule (see Figure 1B). This suggests that conformational exchange and catalysis are coupled. In particular, the crystallographic structures of reduced and oxidized states of *Ec*TPx have shed light on the conformation of the native substates FF and LU. However, no sufficient in-depth dynamic and mechanistic information concerning the conformational transition and protein motions involved is available at the molecular level.

We investigated the FF→LU and LU→FF conformational transitions of *Ec*TPx by computational methods. Conventional MDs allowed us to explore different substates of the conformational landscape. In addition, the different oxidation and protonation states of the catalytic Cys residues involved in the reaction were evaluated. This approach was insufficient to investigate the conformational transition. However, the implementation of aMD allowed an enhancing of conformational sampling. Using this technique, we observed the LU→FF refolding transition suggesting that the process involves high energetic barriers.

It will be very important to study the molecular nature of the barrier, and the degree of coupling between the internal motions, conformational stability of the helical secondary structure elements and the speed of the resolution steps (conformational transition and disulfide bond formation). Our computational results allowed us the study of the LU→FF refolding process, but the FF→LU transition was not observed as such. This conclusion is in agreement with experimental results regarding the secondary structure content (circular dichroism), that indicated that in equilibrium under reducing conditions, an increase in the helical content is observed, which is compatible with the stabilization of the FF ensemble, relative to the LU ensemble, in aqueous solution. Similar behaviors were previously observed for the atypical 2-Cys TPx from *H. pylori* [20] and 2-Cys PrxQ B from *M. tuberculosis* [34].

One of the main questions concerning Prx activity is whether the oxidation/protonation state of the C_P_ might trigger the conformational transition or, on the other hand, there is an inherent equilibrium between FF and LU conformations independent of the oxidation and/or protonation state of C_P_ and C_R_. The available data in the literature is controversial. Some results support the idea that LU conformations can be reached only upon C_P_ oxidation to sulfenic acid [39], while others suggest that the FF/LU equilibrium is independent of the oxidation state of C_P_ [33,40], but in any case, it could be very dependent on each particular Prx. To shed light on this debatable issue, starting from FF and LU conformations, we modeled C_P_ in the thiol, thiolate, sulfenic acid and sulfenate species and subjected to microsecond long cMD simulations. Our MD results suggest that the thiolate species of C_P_ stabilize the FF conformations. This idea is in agreement with the fact that the FF conformations are capable to bind and react with the peroxide substrate [41] since the p*K*a value of C_P_ is typically below 7 [42,43]. However, when aMDs were carried out from the LU conformations we observed that the thiolate form of the Cys residues may establish ionic par interactions that stabilize the LU conformations. On the other hand, when Cys residues were protonated the LU to FF transition occurred. It is interestingly to note that when the disulfide bond of a Prx (in the LU conformations) is reduced by an oxidoreductase (*e.g.* TRX), consecutive thiol/disulfide exchange reactions release C_P_ and the C_R_ both transiently as thiolates [44–46] and then they could receive a proton from the solvent or a neighboring acidic residue. Remarkably, C_P_ and C_R_ as thiols is the protonation state that favors the return to the FF conformations in our simulations. Therefore, we inferred that the deprotonation of the Cys residues can modulate the energetics of the conformational exchange altering the barrier between both conformational ensembles. Protonated states might favour the LU→FF transition, whereas deprotonation might stabilize the existing conformation. A major caveat needs to be given at this point, the reduction reaction and potentially the LU→FF transition happens with the TRX bound to the Prx, such binding (transiently covalent) will importantly alter the electrostatic and solvation environment surrounding the TPx disulfide. TRX is negatively charged at neutral pH (pI = 4.5), the interactions between TPx and TRX are mostly electrostatic and may play a central role in the refolding process.

The conformational exchange between two substates of the native ensemble that allows the formation of the disulfide bond in the active site occurs in other Prxs, including the decameric/dimeric Prx1 and Prx2 in which the reduction of the intersubunit disulfide bonds results in an increase in the secondary structure content [47,48]. An analogue behaviour was previously described in detail for PrxQ, an atypical 2-Cys Prx, which is exclusively found in bacteria, fungi and plants. However, in PrxQ, catalytic Cys is located prototypically in αH2 and typically separated by four residues (C_P_xxxxC_R_), although they can also be functional as 1-Cys [49]. Most PrxQ are monomeric and undergo a significant conformational change upon reduction of the disulfide bond [34]. In the case of *At*PrxQ, thermal unfolding analysis showed that the oxidized enzyme exhibited lower conformational stability than that the reduced enzyme (48 and 53 °C, respectively, [33]). For the oxidized state, it was observed a significant increase in the effective NMR-R_2_ relaxation rates in *At*PrxQ for residues located in the vicinity of the active site [33], compatible with a conformational exchange and localized slow motions in this region of the protein, confirmed by the measurement of *R*_ex_ in CPMG experiments (Carr-Purcell-Meiboom-Gill pulse). This behavior was not observed under reduced conditions [33].

The resolution step (Reaction II, Figure 1), that includes the FF→LU transition, has been shown to be significantly slower than expected for a reaction between a sulfenic acid and a thiol. The rate of resolution is differentially regulated in different Prx and sometimes imposes a severe limitation in the turnover number of Prx [23]. Additionally, changes in the protein sequence can accelerate the resolution step, this has been shown in the cases of *Salmonella typhimurium* AhpC W32F (twofold acceleration, [50]), a mycothiol peroxidase from *Corynebacterium glutamicum* (fivefold acceleration, [51]) and human Prx5 with 22 additional amino acids at the C-terminal (eightfold acceleration, Semelak *et al.* 2019, unpublished results).

Thus, it seems that the conformational exchange between FF and LU substates of the native state may become a rate limiting and is a key step in the catalytic mechanism of Prx regardless the location of the Cys residues: (i) same secondary structure element, (ii) different secondary structure element but located in the same protein subunit, or (iii) Cys residues located in different subunits. It is expected that different Prxs subfamilies exhibit particularities in the FF/LU equilibrium, depending on the protein primary sequence and topology features. Whether or not the molecular characteristics of this step can define important aspects of kinetics is an important question that should be further studied in detail and in a case-by-case basis.

In addition, mutations that destabilize the FF interaction network (particularly F93G and E97G) resulted, not only in increasing probabilities of local unfolding events of the αH3, but also in the modulation of the distance that separates the catalytic Cys residues during a significant portion of the MD simulations, which are critical to understand the catalytic features of the enzyme.

## Materials and methods

### Chemicals

All chemicals reagents were of the purest analytical grade available. Tris(hydroxymethyl)aminomethane (Tris) was from Merck (Darmstadt, HE, DE). Sodium chloride (NaCl) and ethylenediaminetetraacetic acid (EDTA) were from Biopack (Buenos Aires, AR). Isopropyl-β-d-thiogalactopyranoside (IPTG) was from Bio Basic (Amherst, NY, US). Urea was from ICN Biomedicals Inc. (Irvine, CA, US). Dithiothreitol (DTT) and hydrogen peroxide (H_2_O_2_) were from Sigma-Aldrich (St. Louis, MO, US). 4,4’-dithiodipyridine (DTDPy) was from Acros Organics (Hampton, NH, US).

### Expression and Purification of *E. coli* Thiol Peroxidase

The expression vector pPROK1/*tpx* containing the wild-type *E. coli* thiol peroxidase (*Ec*TPx) sequence was a gift of Dr. Derek Parsonage (Wake Forest School of Medicine, Winston-Salem, NC, US). The plasmid was transformed to electrocompetent *E. coli* BL21(DE3) cells and selected using ampicillin 50 μg mL^−1^. The protein was produced in LB medium at 37 °C to OD_600nm_= 0.9 and overexpression was induced by 0.5 mM IPTG for 4 h at 37 °C shaking at 200 rpm. Cells were harvested by centrifugation at 5000 *g* at 4 °C. Pellets were resuspended in 25 mM Tris-HCl, 1 mM EDTA, pH 7.4 (resuspension buffer) on an ice/water bath. Cells were disrupted by pulse-sonication. Cellular debris and insoluble fraction were removed by centrifugation at 7000 *g* at 4 °C. The supernatant fraction containing soluble proteins was loaded onto a sepharose DE52 (Pharmacia Biotech, Uppsala, SE) column equilibrated with 25 mM Tris-HCl, 1 mM EDTA, pH 7.4 at 4 °C and washed with the same buffer. Elution was performed with an increasing gradient up to 1.0 M NaCl in resuspension buffer. Fractions were evaluated by standard SDS-PAGE procedure [52] and UV absorption spectra and those containing *Ec*TPx were pooled and treated with DNase/RNase (50 μg mL^−1^ plus 100 mM Cl_2_Mg during 12 hours at 4 °C), extensively dialyzed against resuspension buffer and then loaded onto a preparative Sephadex G100 (Pharmacia Biotech, Uppsala, SE) size exclusion chromatography (SEC) column (93 × 2.7 cm), previously equilibrated with buffer 10 mM Tris-HCl, 1 mM EDTA, 100 mM NaCl, pH 7.4. Fractions containing *Ec*TPx (>95% pure) were pooled and aliquoted for storage at −20 °C until use.

### Circular Dichroism Spectroscopy Characterization

Changes in secondary and tertiary structure content of wild-type *Ec*TPx were followed by the circular dichroism (CD) signal in the far-UV (195 to 260 nm) using a 1.0 mm path-length cell and in the near-UV (240 to 350 nm) using a 5 mm path-length cell. Measurements were made in a Jasco 810 spectropolarimeter (Jasco Corp., Tokio, JP). *Ec*TPx (8-30 μM final concentration) were prepared in 50 mM Tris-HCl, 1 mM EDTA pH 7.4. Both reduced and oxidized states were obtained incubating the protein samples for 30 minutes at room temperature in the presence of 1.0 mM DTT or 1.0 mM H2O2, respectively. After that, DTT or H2O2 was removed by SEC using a Sephadex G25 matrix (Sigma-Aldrich, St. Louis, MO, US), and the [SH]/[protein] ratios were checked using DTDPy reagent [53] (ε_324nm_ = 21400 M^−1^ cm^−1^). At least ten spectra were acquired at 25 ± 0.1 °C using a Peltier system and a speed scan of 50 nm min^−1^ with a time constant of 0.5 s. The proper blanks spectra were acquired, subtracted to the average protein spectrum, smoothed using a fourth-degree Savitzky-Golay polynomial filter [54] with a 10-point sliding window and expressed to molar ellipticity using the Equation 1:

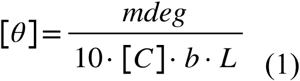

where mdeg is the raw signal in millidegrees, [C] is the protein concentration in molar units, *b* is the number of peptide bonds, and *L* is the path length in centimeters.

### Steady-state Chemical-induced Unfolding Followed by Circular Dichroism Spectroscopy

Equilibrium unfolding experiments were carried out incubating the reduced and oxidized state of *Ec*TPx (25 μM) with increased concentrations of urea up to 7.0 M in 50 mM Tris-HCl, 1 mM EDTA, 100 mM NaCl, pH 7.4 and incubated at room temperature for 5 hours. In the case of the reduced *Ec*TPx, 0.5 mM DTT was added to the buffer to keep reduced conditions. The unfolding process was followed by far-UV CD spectroscopy at 222 nm using a path length cuvette of 0.1 cm as a function of time during at least 200 seconds. All measurements were carried out at 25 ± 0.1 °C. In addition, the far-UV CD spectra at intervals of 1.0 M denaturant were recorder to ensure the loss of structure signatures. A two-state unfolding mechanism (*native* [N]↔*unfolded* [U]) was fitted to the experimental data to calculate thermodynamic parameters [55,56]. The reversibility of the process was checked by dilution in both the reduced and oxidized conditions (data not shown). The final concentration of denaturant was determined by refractive index measurements [57] using a Palm Abbe digital refractometer (*Misco*).

### Oligomeric State in Solution

Size-exclusion chromatography (SEC) was performed in a FPLC system equipped with an UV absorption detector (Jasco Corp., Tokio, JP) and MALLS detector. SEC experiments were carried out using a Superose 12 (GE Healthcare, Chicago, IL, US) column equilibrated in 50 mM Tris-HCl, 100 mM NaCl pH 7.4 prepared in milliQ water and filtered through a 0.22 μm pore size filter. The flow rate was set to 0.3 mL min^−1^ and the injection volume was 100 μL. Protein samples were prepared at 10x (1 mg mL^−1^) to the desired concentration in equilibrium buffer and centrifuged at top speed before injection. In all cases, three independent experiments were performed at room temperature. Data processing was performed using the ASTRA 6.0 software (Wyatt Technology, [58]).

### Computational Characterization of the FF and LU Conformational States

#### Conventional Molecular Dynamics Protocol

The atomic coordinates of the reduced (fully folded, FF conformation) and disulfide bonded (locally unfolded, LU conformation) wild-type *Ec*TPx were retrieved from the RCSB Protein Data Bank under the accession codes 3HVV and 3HVS at 1.75 Å and 1.80 Å of resolution, respectively [13]. Both crystallographic structures were renumbered throughout this work starting from serine 1 in both cases.

The protonation and oxidation state of the peroxidatic Cys (C_P_60) was changed to model different species of interest (see Table 1). The thiol and thiolate protonation state of C_P_ were modeled by the parameters provided in the AMBER package [59]. The parameters for the sulfenic acid species were taken from Defelipe *et al.* [60]. The parameters for the sulfenate species were kindly provided by Dr. Jenner Bonanata (Universidad de la República, Montevideo, Uruguay). The protonation state of the rest of the ionizable residues was set to the corresponding state at neutral pH. The initial coordinates were solvated with the 3-particle TIP3P water model [61] in a cubic box, which extended 10-15 Å further of the nearest protein atom. The set of parameters for alkali halide ions created by Joung and Cheatham [62,63] were used to balance the total charge of the simulation box. A standard minimization protocol was applied to the resulting structures to remove any clashes for 1000 steps of steepest descent followed by 1000 steps of conjugate gradient minimization procedure in the canonical ensemble (NVT) using periodic boundary conditions. The systems were heating up from 100 K to 300 K, using the Berendsen thermostat [61] to suppress fluctuations of the kinetic energy of the system with a relaxation constant of 2 ps and subsequently switched to constant isotropic-isobaric (NpT) conditions to allow the density to equilibrate around ∼1 g cm^−3^ using the Montecarlo barostat. The SHAKE algorithm [64] was applied to all bonds involving hydrogen atoms, then 2 fs time step was settled. An 8 Å cutoff radius for range-limited interactions with the particle mesh *Ewald* summation [65] for long-range electrostatic interactions was used. Harmonic positional restraint of strength 20 kcal mol^−1^ Å^−2^ on Cα atoms was applied during minimization and equilibration and subsequently removed in four successive simulation stages (10, 5, 2 and 0 kcal mol^−1^ Å^−*2*^) of 5 ns each. Unrestrained cMD simulations were performed with the Amber14 [59] suite using the *pmemd*.CUDA engine for the graphics processing unit (GPU) code [66,67] in a GeForce GTX780 (Nvidia) equipped cluster and the *ff14SB* force field [68]. Conventional MD runs were extended for at least 100 ns to obtain the parameters needed and the last restart file was used as a starting point to the accelerated MD simulations.

**Table 1.** Resume of cMD and aMD Simulations Conducted along this Work.

| *EcTPx*<br>conformational<br>substate | Protonation and oxidation<br>states of Cys residues |  | N<sub>atoms</sub><br>protein | N<sub>atoms</sub><br>system | cMD | aMD |  |
| --- | --- | --- | --- | --- | --- | --- | --- |
|  | C<sub>P</sub> | C<sub>R</sub> |  |  |  | λ factor | Simulation<br>steps |
| FF | Thiol | Thiol | 2502 | 29156 | 500 ns | - | - |
|  | Thiolate |  | 2501 | 29147 |  |  |  |
|  | Sulfenic acid |  | 2503 | 29157 |  |  |  |
|  | Sulfenate |  | 2502 | 29148 |  |  |  |
| LU* | Thiol | Thiol | 2502 | 27485 | 500 ns | - | - |
|  | Thiolate |  | 2501 | 27473 |  |  |  |
|  | Sulfenic acid |  | 2503 | 27486 |  |  |  |
|  | Sulfenate |  | 2502 | 27474 |  |  |  |
| LU | Disulfide bonded |  | 2500 | 29188 | 200 ns | - | - |
| FF wild-type | Sulfenic acid | Thiol | 2503 | 27482 | 2000<br>ns | - | - |
| FF R92G |  |  | 2486 | 27473 |  |  |  |
| FF F93G |  |  | 2490 | 27482 |  |  |  |
| FF E97G |  |  | 2495 | 27482 |  |  |  |
| FF | Sulfenic acid | Thiol | 2503 | 35079 | 200 ns | λ=0.3 | 10<sup>8</sup> |
|  | Sulfenic acid | Thiol |  |  |  | λ=1.0 | 2.5x10<sup>8</sup> |
|  | Sulfenic acid | Thiol |  |  |  | λ=2.0 | 4x10<sup>8</sup> |
|  | Sulfenic acid | Thiol |  |  |  | λ=4.0 | 10<sup>9</sup> |
| LU* | Thiol | Thiol | 2502 | 27485 | 100 ns | λ=1.0 | 4 replicas<br>(7.5x10<sup>7</sup>) |
|  | Thiol | Thiolate | 2501 | 35432 |  |  |  |
|  | Thiolate | Thiol | 2501 | 35432 |  |  |  |
|  | Thiolate | Thiolate | 2500 | 35432 |  |  |  |

FF: the fully folded conformational substate. LU, the locally unfolded conformational substate. LU\*: the LU conformational substate lacking the disulfide bond between C<sub>P</sub> and C<sub>R</sub>.

#### Accelerated Molecular Dynamic Simulation Protocol

Accelerated molecular dynamics (aMD) is a biased method that enhances conformational sampling [69], promoting the search for low frequency events in biomolecules by adding a non-negative boost potential ΔV(*r*) to the true potential energy function V(r) when this is lower than a pre-defined energy level E [25,70,71]. Such that, the potential surfaces are raised near the minima and unchanged near the saddle points giving as a result, a decrease in the global potential energy barriers. The biased potential function *V*\*(*r*) is as follows (Equation 2-3):

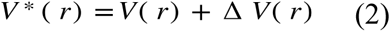

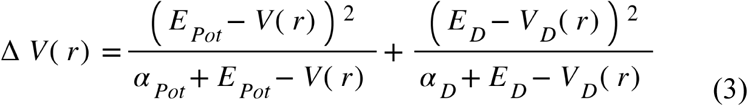

Where *E*_*Pot*_ and *E*_*D*_ are the average potential and dihedral energies obtained by a cMD simulation, V(*r*) and V_*D*_(*r*) are the normal potential and torsion potential, respectively, and α_*Pot*_ and α_*D*_ are the acceleration factors for the potential and dihedral energies, respectively. The dual-boost method was used in all aMD simulations. *E*_*Pot*_, *E*_*D*_, α_*Pot*_ and α_*D*_ are defined as follows:

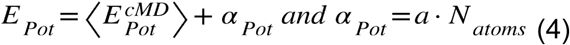

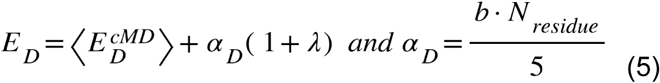

The values of *a* and *b* used in the present work were 0.2 and 3.5 kcal mol^−1^ residue^−1^, respectively. *N*_atoms_ and *N*_residue_ are the total atoms of the system and the number of residues of the protein, respectively. In order to modify the dihedral boost potential, the λ parameter was adjusted as described in Equation 5.

#### General Post-processing Analysis

Time evolution of the protein secondary structure was calculated with the DSSP algorithm [28] of Kabsch and Sander implemented in the *cpptraj* routine [72] of AmberTools15. Solvent accessible surface area (SASA) was calculated using the vmdICE plug-in [73] for VMD 1.9.2. The hydrogen bond (HB) network of the cMD trajectories was calculated using the Python-based HBonanza script [27] and visualized using the provided *Tcl* script for VMD 1.9.2 [74]. The analysis was performed over 2000 frames (200 ns of cMD simulation) setting a donor-acceptor distance of 3.0 Å and hydrogen-donor-acceptor angle of 30 degrees. In order to obtain the most stable HBs along the trajectory, all HBs that were formed less frequently than 50 % of the simulation time were discarded. All protein representations were prepared in VMD 1.9.2 [74].

#### Principal Component Analysis

Principal component analysis (PCA) is one of the widest techniques used for the study of the conformational space of protein dynamics [75,76]. PCA is a quasi-harmonic analysis involving a linear combination to transforms the original dataset of correlated variables into an uncorrelated reduced dimension and the interatomic distance fluctuation covariance matrix *C*_*ij*_ is constructed as follows (Equation 6):

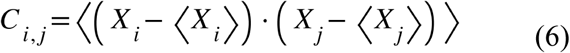

where *x*_*i,j*_ are the Cartesian atomic coordinates of the atom *i,j* and ‹*x*_*i*_› and ‹*x*_*j*_› denote the average coordinates position over the trajectory. After diagonalization of the covariance matrix, a set of eigenvectors and the corresponding eigenvalues are obtained. The orthogonal eigenvectors set describe axes of the maximal variance of the distribution of conformations, and the eigenvalues provide the variance of atomic positional fluctuations contained along each eigenvector. The root-mean-square inner product [77,78] (RMSIP) is a measure for the similarity between the *N* modes of the covariance matrix of two-mode subspaces obtained by PCA and is calculated by Equation 7:

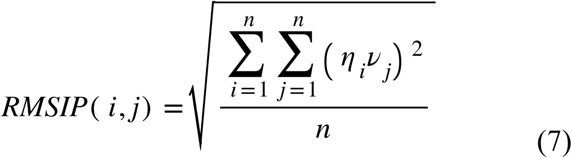

here η_i_ and υ_j_ are the *i*^*th*^ and *j*^*th*^ eigenvectors of each subspace and *n* is the number of eigenvectors evaluated. The two limit values of RMSIP are 0 and 1, where 0 means mutually orthogonal eigenvectors whereas the unity means identical subspaces. In general terms a value of 0.7 is considered an excellent correspondence, a score of 0.5 is fair and below is considered no correlation [77]. The first 10 eigenvectors were used along this work.

## Acknowledgments

This work was supported by grants from the Universidad de Buenos Aires UBACyT N°20020130100468BA, the Agencia Nacional de Promoción Científica y Tecnológica (ANPCyT) PICT N°0983/2013, The Centro Argentino Brasileño de Biotecnología (CABBIO) PICT-CABBIO N°3362/2013 and the Consejo Nacional de Investigaciones Científicas y Técnicas (CONICET) PIP 2010-2015.

## Author Contributions

DSV, AZ, MM, and GFS designed, planned and performed experiments and analyzed data. DSV, AZ, and WAA designed and performed computational simulations and analyzed data. DSV and JS designed and planned all the experiments, analyzed data and wrote the paper. All authors have given approval to the final version of the manuscript.

## Notes

**Conflict of interest:** The authors declare no conflict of interest.

